# Repeated exposure to high-THC *Cannabis* smoke during gestation alters sex ratio, behavior, and amygdala gene expression of Sprague Dawley rat offspring

**DOI:** 10.1101/2023.03.23.533930

**Authors:** Thaisa M. Sandini, Timothy J. Onofrychuk, Andrew J. Roebuck, Austin Hammond, Daniel Udenze, Shahina Hayat, Melissa A. Herdzik, Dan L. McElroy, Spencer N. Orvold, Quentin Greba, Robert B. Laprairie, John G Howland

**Affiliations:** Department of Anatomy, Physiology, and Pharmacology, University of Saskatchewan, Saskatoon, Canada, S7N 5E5; School of Liberal Arts, Yukon University, Whitehorse, YT, Canada, Y1A 5K4; Global Institute for Food Security, Saskatoon, SK, Canada, S7N 4L8; 4Next Generation Sequencing Facility, Saskatoon, SK, Canada, S7N 5E5; Deparment of Oncology, University of Saskatchewan, Saskatoon, Canada, S7N 5E5; College of Pharmacy and Nutrition, University of Saskatchewan, Saskatoon, Canada, S7N 5E5

**Keywords:** prenatal, cannabinoid system, development, behavior

## Abstract

Due to the recent legalization of *Cannabis* in many jurisdictions and the consistent trend of increasing THC content in *Cannabis* products, there is an urgent need to understand the impact of *Cannabis* use during pregnancy on fetal neurodevelopment and behavior. To this end, we repeatedly exposed female Sprague-Dawley rats to *Cannabis* smoke from gestational days 6 to 20 (n=12; Aphria Mohawk; 19.51% THC, <0.07% cannabidiol) or room-air as a control (n=10) using a commercially available system. Maternal reproductive parameters, behavior of the adult offspring, and gene expression in the offspring amygdala were assessed. Body temperature was decreased in dams following smoke exposure and more fecal boli were observed in the chambers before and after smoke exposure in those dams exposed to smoke. Maternal weight gain, food intake, gestational length, litter number, and litter weight were not altered by exposure to *Cannabis* smoke. A significant increase in the male-to-female ratio was noted in the *Cannabis*-exposed litters. In adulthood, both male and female *Cannabis* smoke-exposed offspring explored the inner zone of an open field significantly less than control offspring. Gestational *Cannabis* smoke exposure did not affect behavior on the elevated plus maze test or social interaction test in the offspring. *Cannabis* offspring were better at visual pairwise discrimination and reversal learning tasks conducted in touchscreen-equipped operant conditioning chambers. Analysis of gene expression in the adult amygdala using RNAseq revealed subtle changes in genes related to development, cellular function, and nervous system disease in a subset of the male offspring. These results demonstrate that repeated exposure to high-THC *Cannabis* smoke during gestation alters maternal physiological parameters, sex ratio, and anxiety-like behaviors in the adulthood offspring.

**Significance statement:** *Cannabis* use by pregnant women has increased alongside increased THC content in recent years. As smoking *Cannabis* is the most common method of use, we used a validated model of *Cannabis* smoke exposure to repeatedly expose pregnant rats to combusted high-THC *Cannabis* smoke. Our results show alterations in litter sex ratio, anxiety-like behavior, and decision making in the offspring which may relate to subtle changes in expression of amygdala genes related to development, cellular function, and nervous system disease. Thus, we believe this gestational *Cannabis* exposure model may be useful in delineating long-term effects on the offspring.

## Introduction

Given the recent legalization and decriminalization of *Cannabis* in many jurisdictions around the world, there is an urgent need to better understand its effects in a variety of populations, especially in pregnant women. Indeed, in the United States, up to 7% of pregnant women report using *Cannabis*, which makes it the most prevalent controlled substance used during pregnancy (Volkow et al 2019). *Cannabis* use among pregnant and lactating women is largely motivated by its purported antiemetic properties and for stress and anxiety relief (Skelton et al 2020, Westfall et al 2006). Patterns of *Cannabis* use during pregnancy are variable; however, a significant portion of women continue daily use of *Cannabis* during pregnancy (Ko et al 2015, Metz et al 2022, Pike et al 2021, Satti et al 2022). Population-based studies have also shown long-term effects of maternal *Cannabis* use during pregnancy on offspring development, emotionality, cognition, and brain function (Bara et al 2021, Crume et al 2018, Fried & Smith 2001, Higuera-Matas et al 2015, Moore et al 2022a, Sharapova et al 2018, Smith et al 2006, Smith et al 2016). Cannabinoids readily cross the placenta barrier and modulate endogenous cannabinoid signaling, which is critical for numerous neurodevelopmental processes (Alpar et al 2016, Bara et al 2021, Grant et al 2018, Higuera-Matas et al 2015, Hurd et al 2019, Scheyer et al 2019, Wu et al 2011). However, research on the effects of *Cannabis* on human neonatal outcomes, prenatal development, and long-term behavior are still limited. In addition, the availability of *Cannabis* strains with higher Δ^9^-tetrahydrocannabinol (THC) content has steadily increased in recent years (Smart et al 2017) raising concerns about higher doses of cannabinoids reaching the developing fetus.

Preclinical models are essential to evaluate the effects of gestational *Cannabis* exposure in a more controlled manner, and the information obtained is crucial for a better understanding the underlying neurobiological mechanisms and long-term behavioral consequences (Bara et al 2021, Scheyer et al 2019, Schneider 2009). In rats, gestational exposure to injected cannabinoids (i.e., THC or the synthetic cannabinoid type 1 receptor (CB1R) agonist WIN55,212-2) compromises fetal growth (Natale et al 2020) and normal brain development, leading to cognitive deficits in the offspring (Ferraro et al 2009). Gestational treatment with WIN55,212-2 (0.5 mg/kg, s.c.) from gestational day (GD) 5 to 20, reduces social interaction, ablates endocannabinoid-mediated long-term depression, and heightens excitability of prefrontal cortex pyramidal neurons in male, but not female offspring (Bara et al 2018). Additional studies support these findings, showing the male, but not female, offspring of pregnant dams exposed to THC (2 mg/kg, s.c.; GD5-20) exhibit a hyperdopaminergic phenotype and increased sensitivity to challenge with THC during adolescence (Frau et al 2019, Traccis et al 2021). However, most gestational exposure studies have used maternal injections of cannabinoids, which do not reflect the pharmacokinetics of smoked or ingested cannabinoids or the potential entourage of compounds consumed with whole-plant products (Moore et al 2022b, Russo 2011). Thus, maternal injections have poor face validity when compared to common administration methods use by humans, which include inhalation of *Cannabis* smoke or vaporized extracts and oral consumption. In addition, differences in pharmacokinetics due to the route of administration significantly influence the amount of fetal exposure (Baglot et al 2021, Baglot et al 2022). In this context, the exposure of female rats to vaporized, high-THC *Cannabis* extracts before and during pregnancy altered measures related to anxiety and behavioral flexibility of both male and female offspring (Weimar et al 2020). Thus, use of *Cannabis* exposure paradigms that mimic human consumption patterns are an important goal of recent, ongoing, and future work.

In the present experiments, we chose to assess the effects of repeatedly exposing pregnant rats to the smoke of commercially available dried, high-THC *Cannabis* flowers from GD6-20. The exposure protocol was developed for use in adult rats, and has been shown to result in measurable THC concentrations in plasma, as well as behavioral changes immediately following smoke exposure (Barnard et al 2022, Roebuck et al 2022). During and after *Cannabis* smoke exposure, we quantified an array of maternal and offspring parameters related to acute effects of smoke exposure on the pregnant dams, litter health, and behavior of the male and female offspring in early adulthood. We also assayed the lasting effects of *Cannabis* smoke exposure on gene expression in the amygdala of the offspring using RNAseq. Given the literature reviewed above, we hypothesized that *Cannabis* smoke exposure would alter neonatal development and long-term behavioral phenotypes of male and female adulthood offspring.

## Materials and Methods

### Animals

Sexually naïve female (n=22) and male (n=12) Sprague-Dawley rats (70 days old, Charles River) were pair-housed by sex in a temperature-controlled (21°C) and light-controlled (12h light: 12h dark cycle) vivarium managed by the Lab Animal Safety Unit (LASU) at the University of Saskatchewan. Water and food were available ad libitum in their cages. All procedures were performed during the light phase (0700–1900h) and conducted with the approval of the University of Saskatchewan Animal Research Ethics Board.

After a week of acclimatization to the vivarium, female rats were handled for 3 days (3 min/rat) and were then habituated to the smoke chamber for 20 min/day for 4 days before being bred in the vivarium. The habituation was performed before breeding to reduce stress prior to the early implantation period (GD5). Rats were habituated to the smoke chambers with the pumps off for 2 days, and then with the pumps on for the following 2 days. During the habituation and maternal smoke exposure, dams were placed individually in cages (16 x 25 x 13 (h) cm) in the smoke chambers, and then placed either alone or with a cage in each smoke chamber.

### Breeding

Complete details regarding the breeding protocol are published previously (Sandini et al 2020). One day before breeding began, male rats were split into individual cages. On the day of breeding, 2 female rats (pair-housed) were put in the male cage overnight. The following morning (0800), cells were collected from the vagina of each rat with a sterile P200 pipette tip filled with 50-60 µl of sterile physiological saline. Pregnancies were confirmed by the presence of spermatozoa visualized at the light microscope and this day was considered day 0 of gestation (GD0). After pregnancy was confirmed, rats were singly housed and followed the timeline depicted in Fig. 1. All animals were weighed every 3 days to monitor general health and pregnancy.

**Figure 1.**
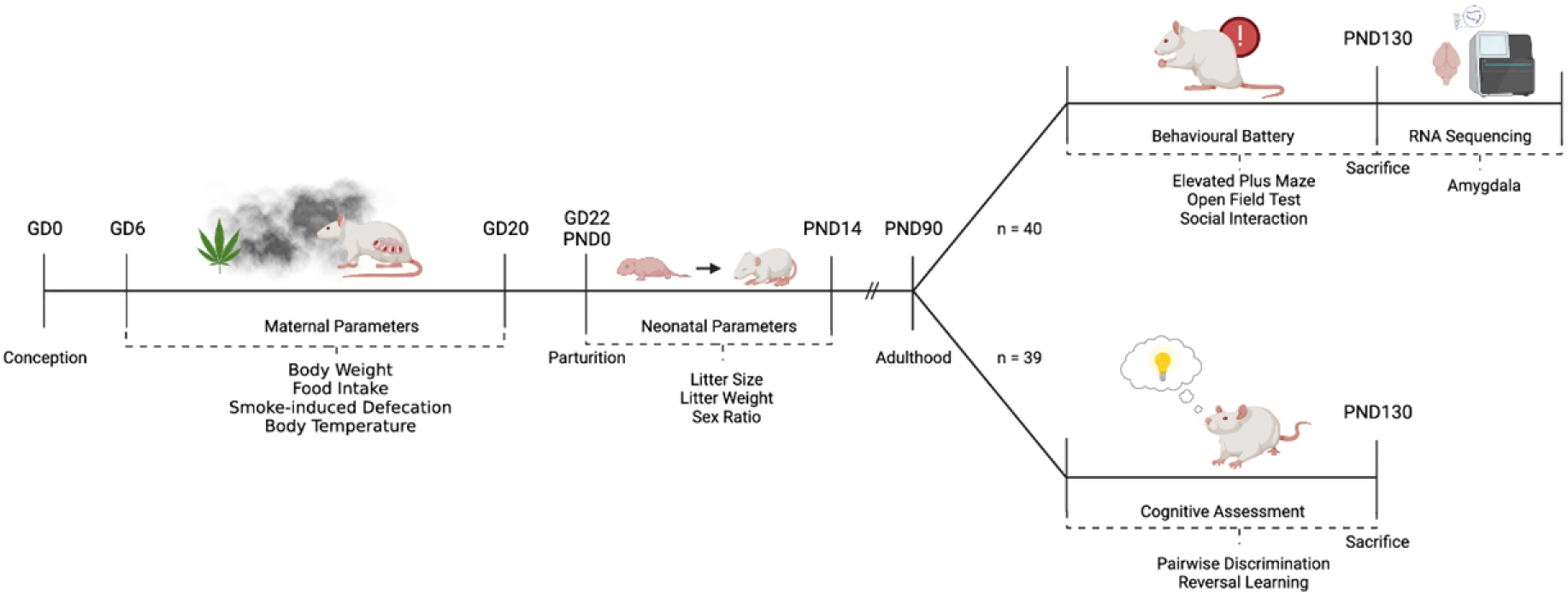
Schematic depicting the flow of dams and their litters through this experiment. Rats were bred in our facility and exposed to either room air or high-THC *Cannabis* smoke from gestational day (GD) 6-20. Litters were assessed for basic parameters during the neonatal period and then left to develop until adulthood (postnatal day (PND) 90). Offspring were then split into 2 cohorts for behavioral and cognitive assessment. Offspring used in the behavioral battery were sacrificed and RNA sequencing was performed on tissue from their amygdala. Figure created with BioRender.com.

### Cannabis

The *Cannabis* strain used in our experiments is called Mohawk (19.51% THC, <0.07% cannabidiol [CBD], https://aphria.ca/product/mohawk/) and was purchased from Aphria (Leamington, ON, Canada). *Cannabis* used in this study was sourced from the same lot (6216) and shipped to our facility in December 2019 and January 2020. Each day, *Cannabis* was freshly prepared by shredding the full flower in a standard coffee grinder (∼5 sec) and weighing it into 200 mg increments. This amount was chosen based on previous work examining the effects of *Cannabis* smoke in rodent models (Barnard et al 2022, Moore et al 2022b, Roebuck et al 2022).

### *Cannabis* smoke exposure system

*Cannabis* smoke exposure was conducted using a validated 4-chamber inhalation system commercially available from La Jolla Alcohol Research, Inc., San Diego, CA used in previous studies (Barnard et al 2022, Roebuck et al 2022). Before each session, *Cannabis* was packed into a ceramic bowl fixed to a metal heating coil which could be heated to combust the product. The coil was sealed with a glass lid and rubber O-ring and the entire assembly could be connected to the atomizer and inhalation system. The inhalation system was composed of 4 identical chambers. Each chamber is airtight and constructed from clear Plexiglas measuring approximately 33 cm (h) x 30.5 cm (w) x 51 cm (l) with an internal volume of ∼50 L. Except during smoke exposure, room air was pumped through the chambers at a flow rate of 10-12 L/min. Air was filtered and exhausted into the facility ventilation system through a fumehood.

On GD6, dams were put into the chambers as described above and after 5 min of acclimatization and equalization of the pressure in all the chambers, the smoke session started. *Cannabis* was combusted over a period of 1 min. To ensure complete combustion, ignition occurred over 5 sec and was repeated three times with a delay of 15 sec between each light. Following combustion, the pumps were stopped for 1 min to allow for exposure. After exposure, pumps were restarted, and the venting process continued for 13 min. The venting was not immediate and significant exposure occurred during this time. After venting was complete, dams were removed and immediately returned to their home cage. Air-control dams were exposed to the same procedure, except that no *Cannabis* was combusted.

### Treatment, maternal, and neonatal parameters

Pregnant rats were split into 2 groups: (i) 200 mg of *Cannabis* combusted and delivered as smoke (n=12) and (ii) room air control (n=10). Exposures occurred once a day for 15 min from GD6-20 (see Figs. 1 and 2). Food intake and body weight were measured every 3 days. Rectal temperatures were also taken prior to and immediately following exposure every 3 days. Furthermore, the number of fecal boli was measured before and after smoke exposure every day. After the last exposure on GD20, the rats were not handled before giving birth on GD21-22. The day of birth was designated postnatal day (PND) 0. On PND 1, litter size, sex ratio, and live birth index were assessed, and the litters were culled to a maximum of 12 pups. When possible, an equal number of males and females were kept. On PND 7 and PND14, litters were weighed again. Due to the COVID19 pandemic, use of the vivarium was severely restricted when offspring entered their second postnatal month in March/April 2020. As a result, litters were culled after weaning to 77 pups for testing in the behavioral battery after PND 90 (Fig. 1).

**Figure 2.**
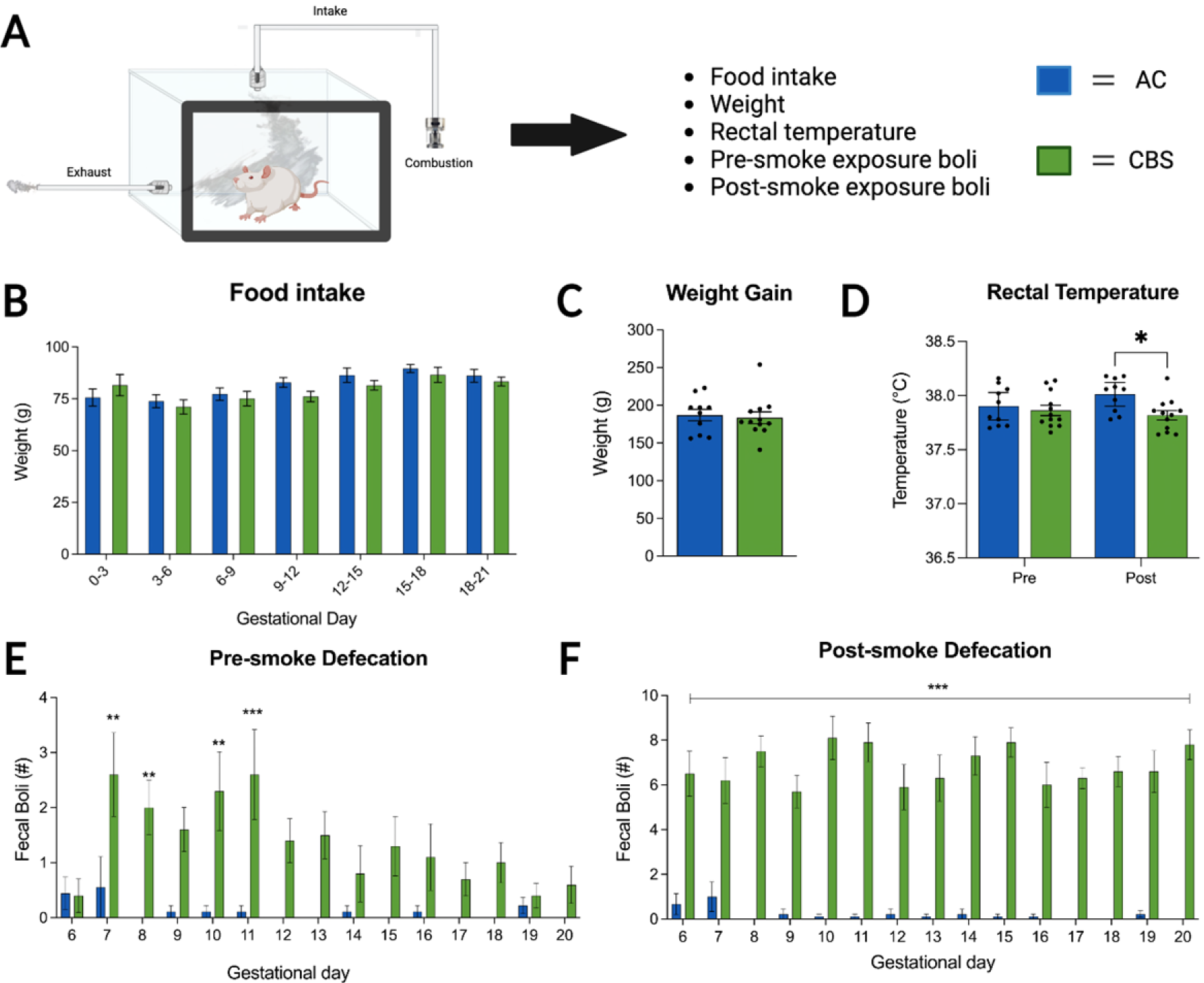
Effects of smoke exposure on the pregnant dams. **A**) Pregnant dams were exposed to either room air or high-THC-containing *Cannabis* smoke from GD6-20. A series of measurements were taken during this phase. *Cannabis* smoke exposure did not affect maternal food intake (**B**) or weight gain (**C**). **D**) Exposure to smoke was associated with a significant decrease in rectal temperature, which was driven by a reduction in temperature post-treatment of the dams treated with high-THC *Cannabis* smoke. Analyses of defecation in the smoke chambers revealed that dams treated with smoke showed increased defecation during the pre-smoke period for several days following the first smoke exposure (**E**). Critically, the groups did not differ before the first smoke administration (GD6). **F**) Dams exposed to smoke consistently defecated more than air controls after the smoke was administered (post-smoke defecation). Asterisks denote significant changes between treatment groups. (*p<0.05, **p<0.01, ***p<0.001) (AC, air control; CBS, *Cannabis* smoke). Figure created with BioRender.com.

### Behavioral testing of the offspring

To test our hypothesis that gestational *Cannabis* smoke exposure causes lasting effects on the behavioral profile of offspring into adulthood, we conducted tests of locomotion, anxiety-like behavior, social behavior and on behavioral flexibility. Testing began around PND90 and was completed by PND130 (adulthood). Offspring were divided into 2 groups: one group (n=40; smoke-exposed male n=11, control male n=9, smoke-exposed female n=11, control female n=9) was first tested in the open field (OF), elevated plus maze (EPM) and social interaction (SI) tests (Fig. 1). A second group (n=37; smoke-exposed male n=10, control male n=7, smoke-exposed female n=11, control female n=9) was tested in the touchscreen-based pairwise discrimination and reversal learning (PD/RL) tasks (Fig. 1). Testing occurred during the light phase (1200-1800) for the OF, EPM, and SI tests, and male rats were tested prior to females. For PD/RL, male rats were trained during the morning (0800-1130) and female rats during the afternoon (1300-1700). Ethanol (40%) was used to clean all behavior testing equipment between rats. Prior to behavioral tests, rats were handled for a minimum of 3 min/day for three consecutive days. Handling included exposure to investigators and emphasized picking up and moving rats until the motions could be carried out with ease, as well as habituation to travel by cart between the animal housing and behavior testing rooms.

#### Open field

The testing apparatus consisted of a circular arena (150 cm in diameter, 45 cm-high walls) made of industrial plastic painted black (Fig. 4A). Rats were brought into the testing room and placed individually in the arena for 15 min. Distance travelled (m) and time spent (sec) in the inner area of the arena was analyzed using Noldus Ethovision XT (Version 6) software.

**Figure 3.**
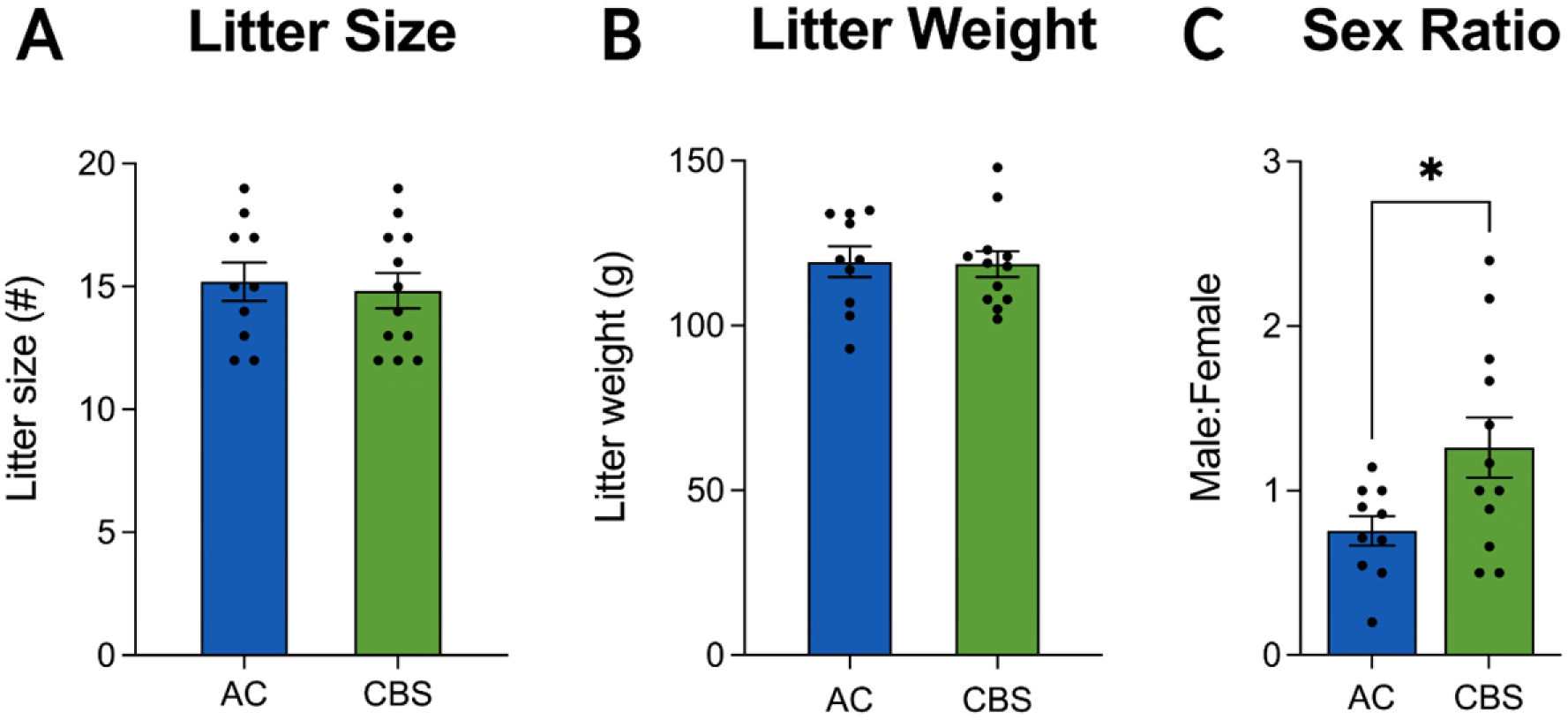
High-THC smoke exposure did not alter the litter size (**A**) or litter weight (**B**), when compared to air controls. **C**) High THC-smoke exposure led to a significant increase in the ratio of male:female pups. Asterisks denote significant changes between treatment groups. (*p<0.05) (AC, air control; CBS, *Cannabis* smoke).

**Figure 4.**
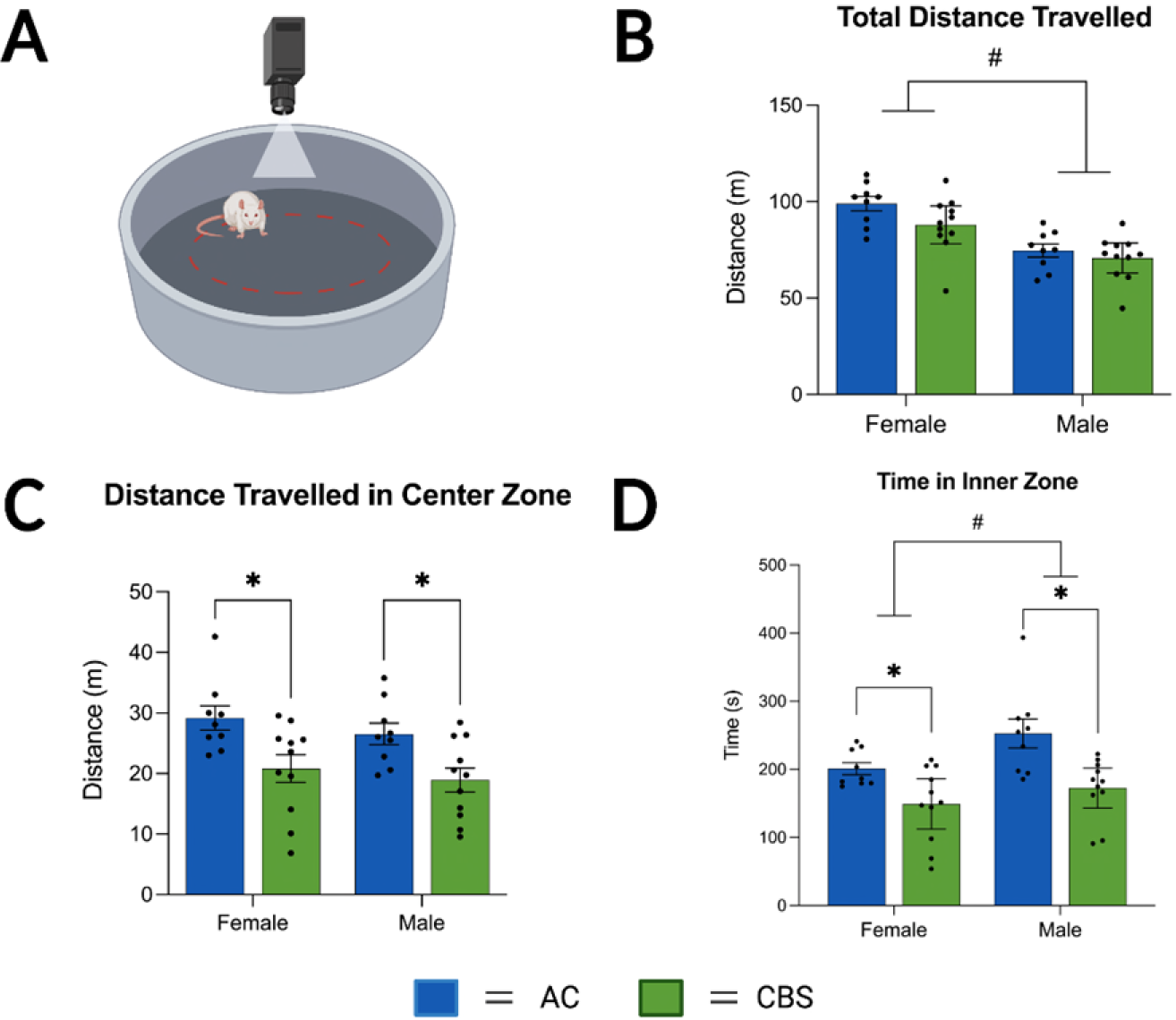
Exploratory and anxiety-like behaviors were assessed in the open field test (**A**). **B**) Female offspring, regardless of treatment, displayed significantly more locomotor activity during the test. The offspring of dams treated with high-THC *Cannabis* smoke traveled less distance (**C**) and spent less time (**D**) in the inner region of the open field. Female offspring, regardless of treatment, also spent less time in the inner zone of the open field. Asterisks denote significant changes between treatment groups (p<0.05). # denotes significant sex differences (p<0.05). Figure created with BioRender.com.

#### Elevated plus maze

The EPM apparatus had 2 open arms and 2 closed arms of equal size (50×10 cm) with 40 cm high walls (Fig. 5A). The EPM was elevated 60 cm above the floor. Rats were placed individually in the central area facing an open arm and were allowed 5 min of free exploration. The number of entries and amount of time in open and closed arms were recorded for each rat. Additionally, rearing, grooming, and head dipping frequencies were evaluated by a researcher blind to treatment.

**Figure 5.**
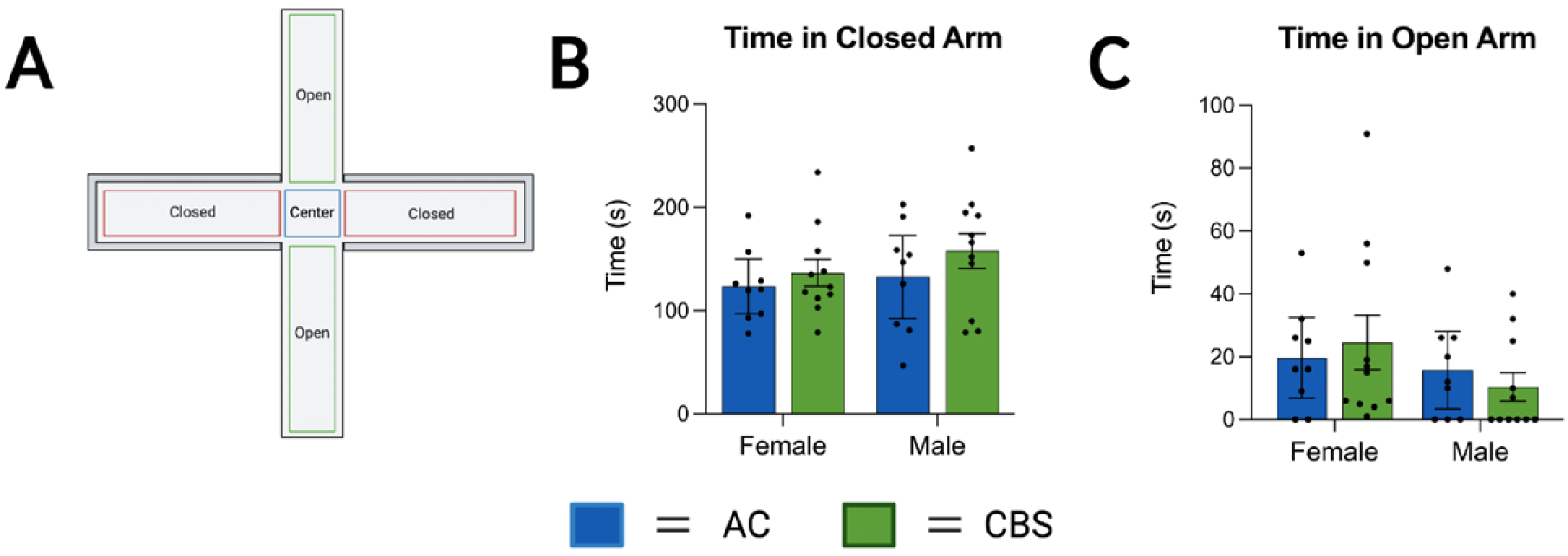
Assessment of the offspring in the elevated plus maze (**A**) revealed no significant differences in the time in the closed arms (**B**) or open arms (**C**). Figure created with BioRender.com.

#### Social interaction

The OF apparatus was used to evaluate social interaction (Fig. 6A). Both test and stranger rats freely explored the circular arena separately for a 10 min habituation period 24 h prior to testing as previously described (Marks et al 2019). During testing, a test rat and a stranger rat of the same treatment and sex were allowed to freely explore the arena for 10 min at the same time. On test day, all test rats were marked on their back with a black marker. Stranger rats were left unmarked. Behavior was scored from videos using stopwatches by a researcher blind to the treatment of the rats. Latency for the first approach (sec) and 20 cm proximity of the test rat were evaluated. The proximity measure included behaviors such as sniffing, nosing, chasing, and passing the stranger rat.

**Figure 6.**
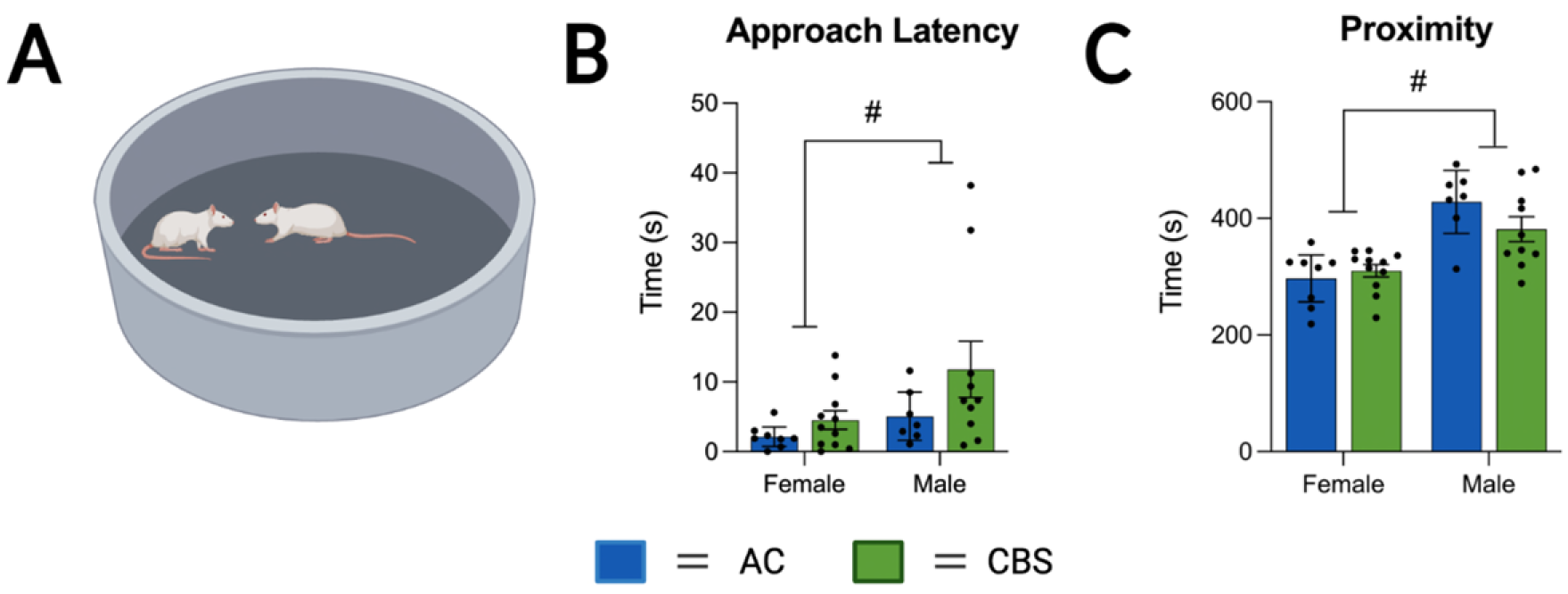
Offspring social interaction behaviors (**A**) were not significantly affected by maternal *Cannabis* exposure. Male offspring approached (**B**) and interacted with 20 cm (**C**) more than female offspring, regardless of treatment. # denotes significant sex differences (p<0.05). Figure created with BioRender.com.

#### Pairwise discrimination and reversal learning (PD/RL) tasks

Discrimination learning and behavioral flexibility was assessed by serial completion of touchscreen-based PD and RL tasks. Seven touchscreen-equipped operant conditioning chambers (Bussey-Saksida Touch Systems, Lafayette Instrument Company, Lafayette, IN, USA) equipped with a trapezoidal inner chamber (30.5cm x 24.1cm x 8.25cm), food reward delivery system (Dustless Precision Pellets, 45 mg, Rodent Purified Diet; BioServ, NJ, USA), overhead camera for live behavioral monitoring, and a touchscreen monitor were used (Fig. 7A). For all training procedures, a black plastic mask and response self was placed in front of the monitor. The mask covered the entirety of the display, except for two identical rectangular cut-outs located in the upper half of the screen where visual stimuli presentation would occur. System functionality checks and cleaning procedures were conducted daily (Barnard et al., 2021).

**Figure 7.**
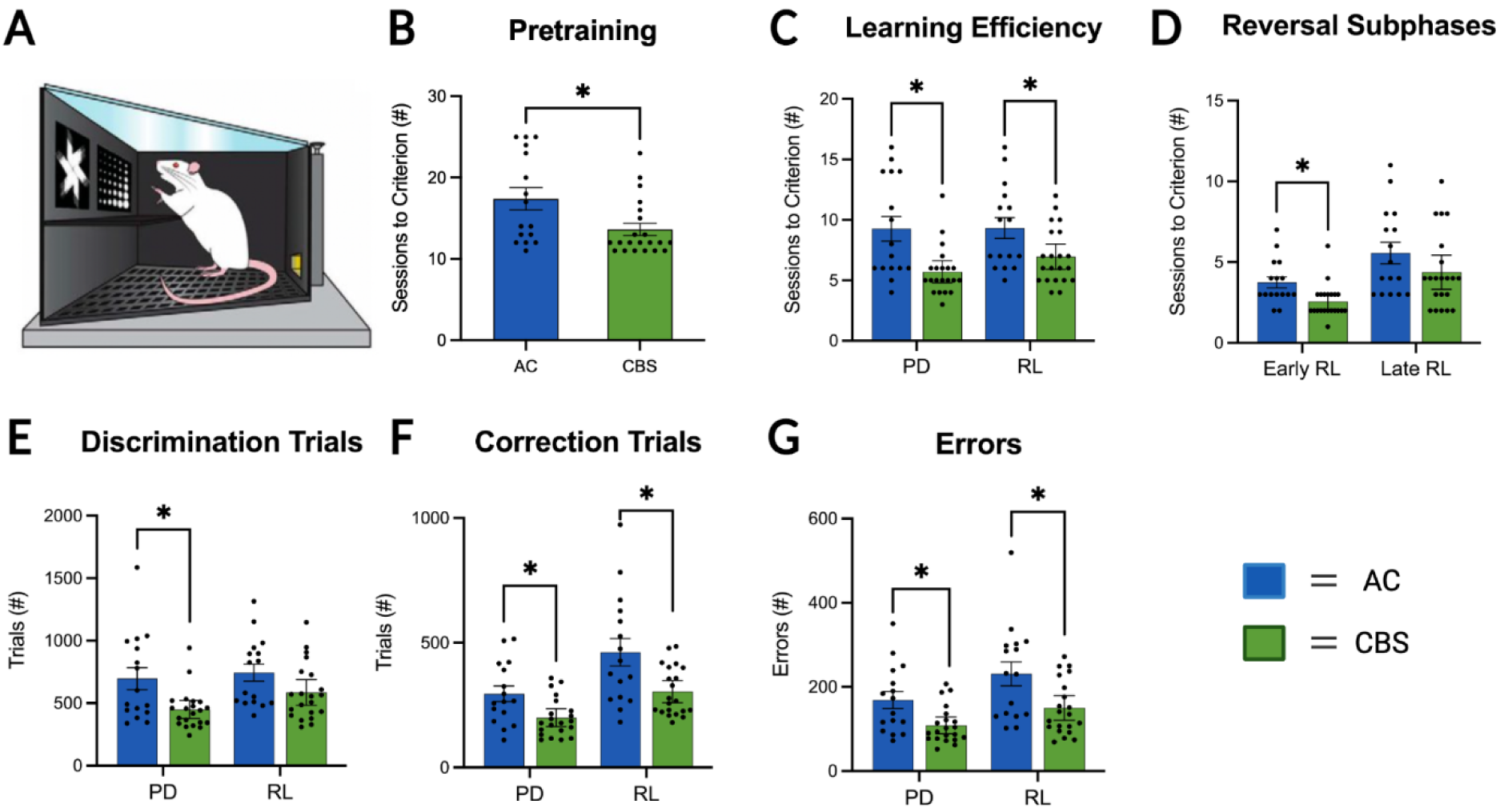
A separate group of offspring were assessed on visual pairwise discrimination (PD) and reversal learning (RL) tests in touchscreen-equipped operant conditioning chambers (**A**). Regardless of sex, offspring of high-THC *Cannabis* smoke-exposed dams required significantly fewer sessions for pretraining (**B**) and to acquire the PD and RL tests (**C**). *Cannabis* offspring required fewer sessions during the early phase of RL (**D**). Significant differences between groups on discrimination trials (**E**), correction trials (**F**), and errors (**G**) are also noted on the figure where appropriate (see text for further details and discussion). Latencies are summarized in Table 1. Asterisks denote significant changes between treatment groups for each task (p<0.05). Figure created with BioRender.com.

**Table 1.** Mean (±SEM) response latencies (sec) for the correct and incorrect trials, as well as reward collection, during the touchscreen testing. No significant differences were observed between the treatment groups for these measures (statistics shown in Supplementary Table 1). (PD, pairwise discrimination; RL, reversal learning; AC, air control; CBS, *Cannabis* smoke.)

|  | PD |  | RL |  |
| --- | --- | --- | --- | --- |
|  | AC | CBS | AC | CBS |
| **Correct** | 5.18±0.8 | 4.16±0.4 | 5.45±0.7 | 4.40±0.4 |
| **Incorrect** | 6.65±1.2 | 4.86±0.6 | 5.09±0.7 | 4.49±0.5 |
| **Reward** | 2.05±0.4 | 1.46±0.1 | 1.49±0.1 | 1.45±0.1 |

Training procedures were conducted as per manufacturer instructions and previous work (Lins et al., 2018; Roebuck et al., 2020). Prior to training, male and female adult offspring were restricted to 85% free-feeding body weight to motivate reward-seeking behavior. Through a series of habituation, pretraining, and training steps, rats were shaped to develop a strong association between visual stimulus selection and a food reward, to initiate stimulus presentation, and to associate incorrection selection with illumination of a bright overhead light. In short, animals were habituated to experimenters, the route of transport, and the testing apparatus over three consecutive days. Next, four stepwise pretraining stages -initial touch, must touch, must initiate, punish incorrect-were completed to develop appropriate stimulus-reward associations and behavioural strategies. Sessions within each stage were 1-hour in duration and restricted to a maximum of 100 selection trials, where advancement between stages was awarded based on established performance criteria. Roebuck and colleagues (2020) provide an in-depth explanation of all habituation and pretraining stages that were used in this study.

Upon completion of pretraining stages, animals advanced to PD, a task measuring visual discrimination learning. Within each selection trial, two distinct black and white images were simultaneously displayed on the touchscreen immediately following trial initiation. One image was always correct, regardless of its spatial location, while the other image was always incorrect. Correct responses were reinforced by a food reward, and incorrect responses were punished by illumination of a bright overhead light and a five-second delay. Incorrect responses were also directly followed by a correction trial. Here, the location of each image was identical to the previously incorrect selection trial, and advancement to a new selection trial could only be achieved by a correct selection. Criterion for advancement in PD was defined as >85% accuracy for 100 selection trials within one hour on two consecutive training sessions. The final phase progression was RL, a test of behavioral flexibility. RL was identical to PD, except the correct image is reversed; said differently, selection of the previously incorrect image was rewarded, and selection of the previously correct image resulted in a five-second delay and a subsequent correction trial. Criterion for completion was defined as >85% accuracy across 100 selection trials within one hour on two consecutive training sessions. Two male rats (1 from each treatment group) failed to complete RL and as a result, their data were not included in the analysis.

### Statistical analysis – behavioral data

Data were analyzed using GraphPad Prism 8.0.1 software. All figures show means with the error bars showing the standard error of the mean (SEM). Prior to analysis, data were tested for normality of distribution using the Shapiro-Wilk test. Unpaired t test was used to analyze gestational length, litter size and weight and male/female ratio. Two-way ANOVA (followed by Bonferroni’s multiple comparisons test) with factors of Treatment (air control vs. *Cannabis* smoke) and Time (gestational days) were used to evaluated maternal gain weight, food intake, body temperature and pre- and post-boli smoke exposure. For the offspring behavioral testing, two-way ANOVA (followed by Bonferroni’s multiple comparisons test) with factors of Treatment (air control and *Cannabis* smoke) and Sex (male, female) was used when data were parametric. Non-parametric Mann Whitney tests were used when data were non-parametric. *P* values of < or = 0.05 were considered significant.

### Gene expression in the offspring amygdala

#### Sample collection

After behavioral testing, the offspring from the OF, EPM, and SI cohort (n=40) were anesthetized with isoflurane and decapitated. Amygdala tissue samples were quickly dissected, and flash frozen in liquid nitrogen for gene expression analysis.

#### RNA collection and library preparation

Total RNA was extracted from the amygdala of adult rats using the MagMAX mirVana Total RNA Isolation kit (ThermoFisher). RNA quality was assessed using the Qubit RNA BR Assay (ThermoFisher) and RNA Screentape (Agilent). Sequencing libraries were constructed from 300 ng of RNA per sample using the TruSeq Stranded mRNA Library Prep kit (Illumina).

#### RNA sequencing

Sequencing libraries were evaluated using the Qubit dsDNA BR Assay (ThermoFisher) and a D1000 Screentape (Agilent). The barcoded libraries were pooled equimolar and 75 bp paired-end reads were generated on a NextSeq 550 instrument (Illumina).

#### Data processing

The reads were extracted from each run using bcl2fastq (version 2.19.0.316). Sequencing adapters and low-quality bases were trimmed using fastp (Chen et al 2018) with default settings. The reads were aligned to the *Rattus norvegicus* reference genome (GCA_000001895.4) using subjunc (from subread version 2.0.1) and gene-level expression quantified using htseq-count from the HTSeq framework (version 0.11.3) with default settings except for: “--nonunique all, stranded=reverse” (Anders et al 2015).

#### Differential Gene Expression

Differential expression between exposed and unexposed rats was assessed using Autonomics (version 1.3.0) in RStudio (version 4.1.2). Using Autonomics pipeline, RNA expression counts were log transformed and fitted to the “∼ 0 + subgroup” model using the limma package. Significant genes for the contrast between rats exposed to *Cannabis* smoke during gestion (Treated) and unexposed (Control) rats were extracted using Benjamini-Hochberg adjusted *p* value (FDR < 0.05) and absolute fold-change of 1.5. Signaling pathways, biological and disease functions, and regulatory network interactions associated with differentially expressed genes were identified using QIAGEN Ingenuity Pathway Analysis (IPA) (QIAGEN Inc.). Genes with absolute fold change greater than 1.5 were included and molecules and relationships were filtered for “tissues/cell lines=amygdala, brain, CNS cell lines, or fibroblast cell lines” were included. Raw data related to these analyses have been posted on the NCBI repository under Bioproject Accession number PRJNA886744.

## Results

### Prenatal *Cannabis* smoke exposure alters maternal responsivity to the smoke chambers and offspring sex ratio

Pregnant rats were exposed to *Cannabis* smoke once daily from GD6-20 and a variety of measures were taken from dams and offspring at birth (Fig. 1, 2A). Gestational *Cannabis* exposure did not affect maternal food intake (Fig. 2B) or weight gain (Fig. 2C) during pregnancy (statistics shown in Supplementary Table 1). However, rectal temperature was significantly affected by treatment, with rats in the *Cannabis* smoke group showing significantly lower temperatures after treatment (Fig. 2D; Treatment by Time interaction F(_1,20_) = 10.52, p = 0.0041; posthoc p < 0.05), with no main effect of Time (F(_1,20_) = 1.85, p = 0.19) or Treatment (F(_1,20_) = 3.21, p = 0.09).

When compared to air controls, rats exposed to *Cannabis* smoke from GD6-20 defecated more in the smoke chambers during pre- and post-smoke periods. Analysis of pre-smoke boli showed significant main effects of Treatment (Fig. 2E; F(_1,255_) = 71.47, p < 0.0001) and Gestational Day (F(_14,255_) = 1.94, p = 0.02), and a significant interaction (F(_14,255_) = 1.72, p = 0.05). Post-hoc analyses indicated that smoke-exposed dams produced significantly more boli on GD7, GD8, GD10, and GD11 than air controls. During the post-smoke period, *Cannabis*-exposed dams produced significantly more boli than air controls regardless of GD (Fig. 2F; main effect of Treatment: F(_1,255_) = 755.0, p < 0.0001, with no effect of Gestational Day (F(_14,255_) = 0.69, p = 0.78) or an interaction (F (_14,255_) = 1.03, p = 0.42).

Dams were allowed to give birth naturally and litters were assessed on PND1. No differences were observed in the number of total pups born (Fig. 3A) or total litter weight on PND1 (Fig. 3B). However, a significant increase in the male:female ratio was observed in litters exposed to *Cannabis* smoke (*t*_20_ = 2.34, p = 0.03). All litters were culled to 12 pups per dam. The average litter weight on PND7 (Air control = 230.6±5.2; *Cannabis-*exposed = 231.6±5.9) and PND14 (Air control = 406.2±8.0; *Cannabis-*exposed = 404.7±10.7) were not significantly altered by *Cannabis* smoke exposure (statistics shown in Supplementary Table 1).

### Gestational *Cannabis* smoke exposure alters activity in the OF, but not EPM or SI, tests

Distance travelled (m) within the entire circular arena (Fig. 4A) was analyzed to examine general locomotor activity in the OF (Fig. 4B). Gestational *Cannabis* smoke exposure did not alter the total distance traveled (Fig. 3A; F(_1,36_) = 3.69, p = 0.06); however, there was a significant effect of Sex (F(_1,36_) = 29.37, p = 0.0001) without an interaction (F(_1,36_) = 0.83, p = 0.36). Inspection of the data revealed female offspring travelled further than their male siblings, regardless of treatment. To assess anxiety-like behavior, we quantified distance travelled (Fig. 4C) and time spent (Fig. 4D) in the inner zone of the open field. Analyses of these data showed that the *Cannabis* smoke-exposed offspring travelled less and spent less time in the inner zone (main effect of Treatment for distance (F(_1,36_) = 15.02, p < 0.001) and time (F(_1,36_) = 17.80, p < 0.001)) of the OF. There was also a main effect of Sex for time in the inner zone (F(_1,36_) = 5.74, p = 0.02), with females spending less time in the inner zone than males, regardless of treatment (Fig. 4D). All other main effects and interactions were not significant (statistics shown in Supplementary Table 1).

The EPM test (Fig. 5A) was used as another index to measure anxiety-like behavior following gestational *Cannabis* smoke exposure in the same offspring as were used for the OFT. Two-way ANOVAs indicated no main effects of Treatment, Sex, or interactions when time spent exploring the closed and open arms (Fig. 5) was considered (statistics shown in Supplementary Table 1). Also, no differences were noted in the number of open and closed arm entries, rears, time spent grooming, or head dips were analyzed (all *p*’s>0.05; data not shown).

In the SI test (Fig. 6A), latency (sec) to first approach (Fig. 6B) and proximity (Fig. 6C) were evaluated. Two-way ANOVA indicated a main effect of Sex (F(_1,32_) = 4.14, p = 0.050), but not Treatment (F(_1,32_) = 3.29, p = 0.079), nor an interaction (F(_1,32_) = 0.75, p = 0.39), for latency to first approach. When we analyzed the time spent within a 20 cm proximity, two-way ANOVA indicated a main effect on Sex (F(_1,32_) = 31.62, p < 0.0001), with no Treatment effect (F(_1,32_) = 0.86, p = 0.36) or interaction (F(_1,32_) = 2.75, = 0.11). Inspection of the data revealed that male offspring approached each other and spent significantly more time within 20 cm of the stranger rat, regardless of treatment.

### Effects of prenatal *Cannabis* smoke exposure on pairwise discrimination and reversal learning task (PD/RL)

A separate cohort of male and female offspring were tested for PD and RL (Fig. 7A). Regardless of Sex, rats from dams treated with *Cannabis* smoke (n=16) completed the pretraining phases of task acquisition faster than those from control dams (n=21; Fig. 7B; Mann Whitney test p = 0.011). When sessions to complete PD and RL were considered (Fig. 7C), analyses revealed *Cannabis-*treated offspring learned the PD and RL rules in significantly fewer sessions than air controls (Mann Whitney tests, p=0.002 for PD, p=0.02 for RL). Analysis of sessions to complete early RL (i.e., all sessions before rats achieve greater than 50% correct on the reversal) and late RL (i.e., all sessions after rats achieve 50% correct on the reversal) revealed a significant difference in early, but not late, RL (Fig. 7D; p=0.002 Mann Whitney). It is noteworthy that *Cannabis* offspring were also faster than controls during the late RL phase, although results were not significant.

A more detailed analysis of performance of the offspring during the PD and RL tests revealed subtle differences between treatment groups. When the number of discrimination trials were analyzed (Fig. 7E), *Cannabis*-exposed offspring required significantly fewer trials to reach criterion in the PD, but not RL, test (Mann Whitney, p=0.015). Analyses of correction trials (Fig. 7F) and errors (Fig. 7G) revealed that offspring exposed to *Cannabis* smoke completed fewer correction trials and made fewer errors than control offspring in both PD and RL (all Mann Whitney p values ≤ 0.02). Latencies for correct responses, incorrect responses, and reward collection for PD and RL are shown in Table 1. No significant differences were seen between the treatment groups for any of these measures (statistics shown in Supplementary Table 1).

### Prenatal *Cannabis* exposure altered amygdala gene expression in the adulthood offspring

Next, whole-genome transcriptomics of amygdala tissues of adult rat offspring exposed to *Cannabis* smoke during gestation or control conditions were studied using RNA-seq. Principal component analysis (PCA) of the expression data showed no obvious clusters corresponding to treatment group within the female samples, and as a result they were excluded from further analysis. Furthermore, in the male cohorts, two control male samples and three treated male samples were observed to cluster with samples from the opposite condition and were also excluded from further analyses. Full transcriptome analysis of the remaining 15 samples revealed a clustering effect in the male offspring exposed to Cannabis smoke (n=7) and controls (n=8; Fig. 8A). Male offspring exposed to *Cannabis* during gestation had profound transcriptional changes with 1746 differentially regulated genes of which 876 were upregulated and 870 were downregulated (FDR-adjusted P < 0.05; absolute fold change (FC) > 1.5) (Fig. 8B, Supplementary Table 2A). Among the top 40 affected genes were 23 genes linked to neuronal functions, behavior, or neurodevelopmental disorders (GeneIDs are detailed in Supplementary Table 2, sheet A): *Ap2a1* (Vrahatis et al 2020), *Cers1* (Godeiro Junior et al 2018), *Gdf1* (Lee 1991), *Hspbp1* (Jing et al 2021), *Fip1l1* (Tennenbaum et al 2021), *Msi2* (Luan et al 2016), *Ssbp1* (Meunier et al 2021), *Tmx3* (Chao et al 2010), *Tmem179* (Carpanini et al 2017), *Usp20* (Jolly et al 2022), *Hspb11* (Shields et al 2019), *Ints1* (Zhang et al 2020), *P4htm* (Kraatari-Tiri et al 2022), *Atp13a2* (Pan & Yue 2014), *Ccdc120* (Abidi et al 2016), *Dpp3* (Ren et al 2021), *Fam69b* (Dudkiewicz et al 2013), *Fbxo31* (Vadhvani et al 2013), *Prr7* (Lee et al 2018), *Rabac1* (Fenster et al 2000), *Neurl4* (Imai et al 2015), *Slc35a2* (Vals et al 2019), and *Smpd1* (Alcalay et al 2019).

**Figure 8.**
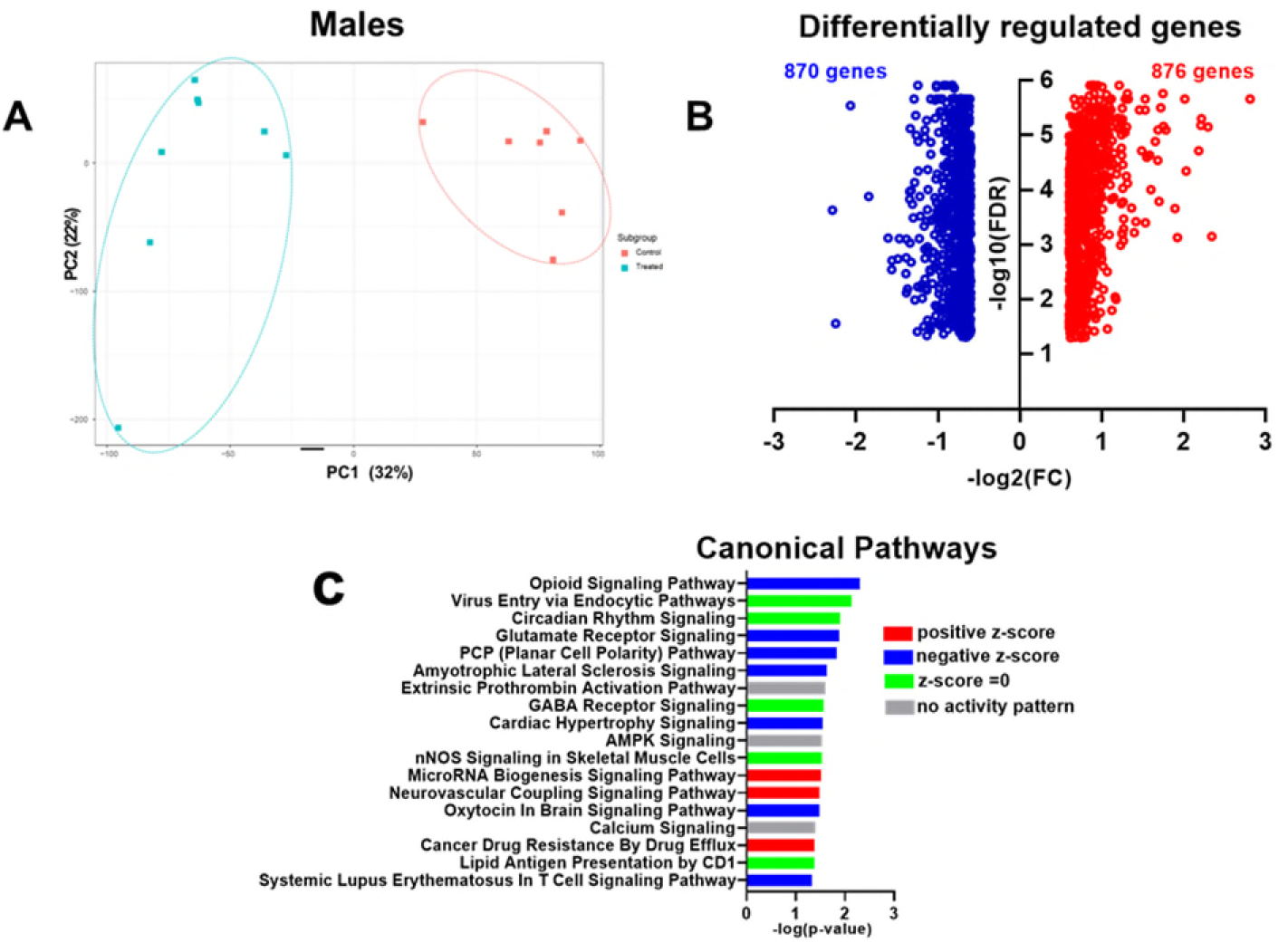
Transcriptional responses in the amygdala of male rat offspring exposed to high-THC *Cannabis* smoke during gestation. **A**) Principal-component analysis (PCA) of RNA-seq data of male animals. **B**) Volcano plots display significantly affected genes (FDR < 0.05). Plots of the upregulated (red) and downregulated (blue) genes. **C**) Significantly enriched canonical pathways. Positive z-score indicates predicted activation; negative z-score indicated predicted inhibition. z-score of zero indicate no clear direction of activity. No activity pattern indicates not sufficient information in the Ingenuity Knowledge Base. Raw data are in Supplementary Table 2A and B.

Accordingly, Ingenuity Pathway Analysis showed significant effects in male offspring exposed to *Cannabis* smoke during gestation with alterations in 18 canonical pathways by applying −log (P-value) >1.3 threshold (Fig. 8C; Supplementary Table 2B). Among the top 18 most enriched pathways were 7 pathways involved in Neurotransmitters and Other Nervous System Signaling: Opioid Signaling, Circadian Rhythm Signaling, Glutamate Receptor Signaling, GABA Receptor Signaling, Amyotrophic Lateral Sclerosis Signaling, Neurovascular Coupling Signaling, and Oxytocin in Brain Signaling (Fig. 8C).

Functional set enrichment analysis for disease and biological processes also showed significant effects in amygdala of male offspring exposed to *Cannabis* during gestation (Fig. 9A; Supplementary Table 2C, D and E). Specifically, genes with altered expression in the amygdala of exposed rats were predicted to be negatively enriched for processes related to Nervous System Development, Cellular Functions, and Maintenance and Neurological Disease.

**Figure 9.**
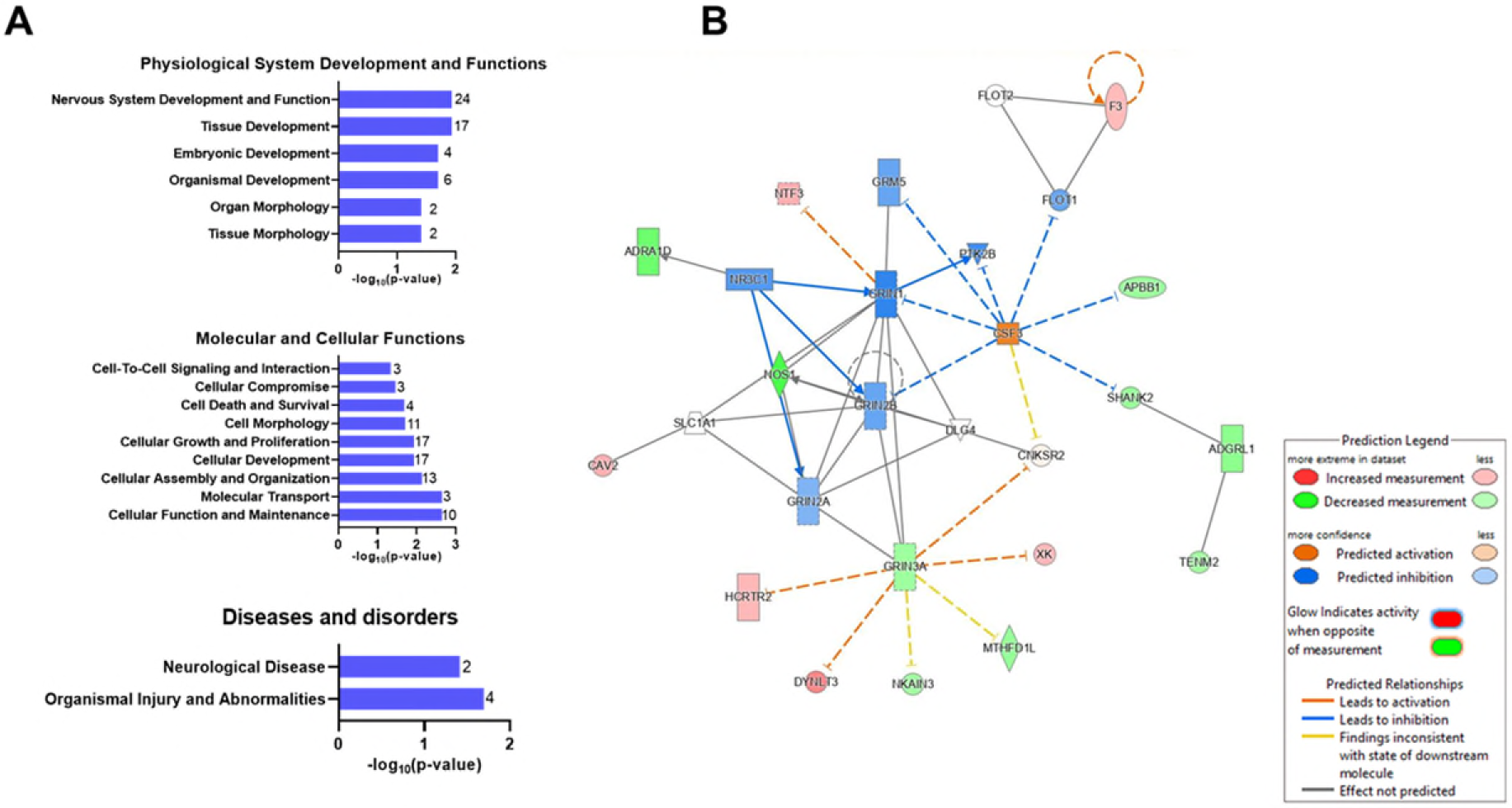
Gene-ontology analysis of RNA-seq analysis from amygdala samples detailed in Figure 8. **A**) Gene enrichment analysis of differentially expressed genes for biological and disease function identified perturbed pathways in physiological system development and functions, molecular and cellular, disease and disorders. Data are represented as -log_10_ of the lowest p-value of an associated pathway. Values at the end of each bar represent the number of differentially expressed genes in each gene-ontology category. **B**) Gene interaction network map. This network consists of the top 2 ranked networks of differentially expressed genes linked to Neurological Disease, Organismal Injuries and Abnormalities, and Psychological Disorders.

Next, we performed network interaction analysis between differentially regulated targets in rats exposed to *Cannabis* during gestation. The top 5 ranked networks were networks of genes involved in Cellular Development and Functions, Nervous System Development and Functions, Neurological Disease, Behavior, Organismal Injuries and Abnormalities, and Psychological Disorders. The top 2 interaction network maps, which are involved in Neurological Disease, Organismal Injuries and Abnormalities, and Psychological Disorders, have 27 nodes including 15 focus genes from whole genome transcriptomic targets— *Adgrl1*, *Adraid1*, *Apbb1*, *Cav2*, *Cnksr2*, *Csf3*, *Dlg4*, *Dynlt3*, *F3*, *Flot1*, *Flot2*, *Grin1*, *Grin2a*, *Grin2b*, *Grin3a*, *Grm5*, *Hcrtr2*, *Mthfd1l*, *Nkain3*, *Nos1*, *Nr3c1*, *Ntf3*, *Ptk2b*, *Shank2*, *Slc1a1*, *Tenm2*, and *Xk* (Fig. 9B; Supplementary Table 2F).

## Discussion

To the best of our knowledge, the present study is the first to examine the effects of commercially available high-THC *Cannabis* smoke administration on rats during pregnancy and their offspring. We show that smoke exposure increases defecation in pregnant rats during the exposure period and significantly reduces their temperature following exposure (Fig. 2). Maternal weight gain and food intake were not altered by the smoke exposure protocol (Fig. 2). Litter number and size were not affected by maternal smoke treatment, although the sex ratio of male to female offspring was significantly increased (Fig. 3). In adulthood, offspring of dams exposed to *Cannabis* smoke during gestation showed evidence of anxiety-like behavior in the open field test (Fig. 4), while treatment effects in the elevated plus maze (Fig. 5) and social interaction tests (Fig. 6) were not observed. Offspring of rats exposed to *Cannabis* during gestation showed facilitated learning of a visual paired discrimination and subsequent reversal in touchscreen-equipped operant conditioning chambers (Fig. 7). Analyses of amygdala gene expression demonstrated altered expression of genes related to nervous system development and function in a subset of male, but not female, offspring (Figs. 8 and 9). Taken together, these findings suggest that exposure to high THC *Cannabis* smoke alters neural circuits involved in emotional and cognitive behaviors.

### High THC *Cannabis* smoke exposure produced acute changes in physiology of the pregnant dams without dramatic changes in fetal and neonatal parameters

Our maternal *Cannabis* smoke exposure paradigm involved a single daily exposure from gestational day 6 to 20, which corresponds to the first and second trimesters of human pregnancy. *Cannabis* use is more prevalent during these stages of pregnancy as undesirable symptoms such as nausea are more prevalent (Volkow et al 2019). In addition, we did not treat the dams during the earliest stages of pregnancy given concerns regarding the effects of cannabinoid exposure on implantation and fetal reabsorptions. Daily exposure to high THC *Cannabis* smoke did not affect the amount of food consumed nor maternal weight gain during gestation, thus minimizing concerns related to potential malnutrition on fetal development. However, *Cannabis* smoke exposure significantly decreased rectal temperature, consistent with effects seen in some studies following THC vapor exposure in female rats (Javadi-Paydar et al 2018, Nguyen et al 2016), but not others with pregnant rats (Breit et al 2020). Tolerance to the hypothermic effects of THC vapor exposure have also been noted following twice daily exposure for 14 days (Nguyen et al 2020). Given the differences in the effects of inhaled vapor or smoke containing THC, analyses of blood levels of THC and its metabolites would help in understanding these differing effects. While we did not measure blood levels of cannabinoids following smoke exposure in this study, acute exposure to the smoke from 300 mg of high-THC containing smoke results in plasma levels of 10-20 ng/ml after 30 min (Barnard et al 2022) and drops to < 5 ng/ml 75 min after exposure (Roebuck et al 2022). While it is difficult to extrapolate the plasma levels following acute exposure to 300 mg of *Cannabis* flower in adult rats to repeated exposure of pregnant rats to 200 mg of *Cannabis* flower, our exposure protocol likely produced peak plasma levels that approach the low end of typical blood levels detected in humans (Newmeyer et al 2016).

Exposure to *Cannabis* smoke using our protocol did not affect litter size or litter weight during the first 2 postnatal weeks. Others have confirmed that inhaled THC exposure during pregnancy in rats does not affect offspring number or weight around the time of parturition (Baglot et al 2022, Breit et al 2020), although reduced pup weight has been noted in treated offspring in protocols that exposed rat dams to vaporized THC twice daily during mating and pregnancy (Weimar et al 2020) and a study of *Cannabis* smoke exposure during gestation in mice (Benevenuto et al 2017). Thus, both timing of smoke exposure and rodent species could be factors in determining the effects of *Cannabis* smoke on birthweight. In humans, findings related to *Cannabis* use during pregnancy and fetal/neonatal growth are mixed (Grant et al 2018, Nashed et al 2020). Therefore, further research in this area is warranted.

Interestingly, exposure to *Cannabis* smoke significantly increased the male:female sex ratio of the litters. Some previous studies have also reported this effect in mice (Benevenuto et al 2017) and rats (Hutchings et al 1987) following exposure to *Cannabis* smoke or THC. As administration of *Cannabis* smoke was initiated on GD6 in the present study, mechanisms related to the viability of female fetuses following implantation may be more sensitive to the effects of THC, or other constituents of *Cannabis* smoke as administered in the present study.

### Offspring exposed to high-THC *Cannabis* smoke during gestation have altered anxiety-like behavior and discrimination learning in adulthood

A subset of adult offspring was tested in the OF, EPM, and SI tests in adulthood to assess locomotor responses, anxiety-like behavior, and social behavior. Total distance traveled in the OFT was not altered in *Cannabis* smoke-exposed adult offspring, which is generally consistent with previous studies (Campolongo et al 2011), and those that show changes in locomotor activity following gestational *Cannabis* exposure only after pharmacological challenge with drugs such as amphetamine (Silva et al 2012) and THC (Frau et al 2019). Offspring exposed to *Cannabis* smoke showed reduced distance and time in the inner zone of the OF, measures often used as indices of anxiety-like behavior in rodents. In the EPM, open arm time, a measure related to anxiety-like behavior, did not differ between the treatment groups. While it is unclear why results differed between these two tests, it is noteworthy that several rats in each group failed to enter the open arms in the EPM. Thus, the room environment for that test may have induced a relatively high baseline level of anxiety in the rats and obscured group differences (i.e., due to a floor effect). Other studies, where open arm exploration was higher in control rats, have revealed an anxiogenic profile in adult rat offspring exposed to THC during gestation (Trezza et al 2008, Weimar et al 2020); but see also (Bara et al 2018). Data from the SI test showed relatively subtle differences between treatments for initial approach latency and proximity that failed to reach significance, particularly for the male offspring. Regardless of treatment, male rats in general spent significantly more time near each other during the test than their female counterparts. Previous studies of the effects of gestational cannabinoid exposure on social interaction in rats are inconsistent with some showing changes in juvenile (Weimar et al 2020) and adult offspring (Bara et al 2018), while others show no effect of gestational cannabinoid exposure on social interaction in the offspring (Manduca et al 2020, Traccis et al 2021).

Behavioral flexibility is an executive function essential for optimizing behavioural responses to a dynamic environment and is disrupted in many psychiatric and neurodevelopmental disorders (Hurtubise & Howland 2017, Soltani & Izquierdo 2019, Uddin 2021). Existing preclinical work shows that behavioral flexibility is altered in prenatal exposure models, including maternal immune activation, autism spectrum disorder, and fetal alcohol spectrum disorder (Ballendine et al 2015, Lins et al 2018, Marquardt et al 2014, McKinnell et al 2021, Zhang et al 2012). Here, we utilized a touchscreen-based visual discrimination (PD) and reversal learning (RL) paradigm to assess executive function in *Cannabis*-exposed offspring. *Cannabis*-exposed offspring required fewer training sessions to complete PD and RL, suggesting a *Cannabis* smoke-mediated facilitation effect. This facilitation is reflected in several related, but dissociable, performance measures as *Cannabis*-exposed offspring completed these tasks with fewer discrimination trials, correction trials, and errors. Notably, a subset of air control offspring in the present sample performed worse on both PD and RL relative to previous studies, potentially driving the observed facilitation effect (Bryce & Howland 2015, Lins et al 2018). In contrast to the present findings, Weimer and colleagues (2020) found that exposure of rat dams to high-THC *Cannabis* vapor before and during pregnancy marginally increased the number of trials offspring required to reach criterion on a lever-based visual cue discrimination, but not reversal learning, task. Hernandez and colleagues (2021) report that adult rats exposed to *Cannabis* smoke in adolescence exhibit enhanced performance on a delayed response working memory task; however, generalizability of enhancement to the present study is limited by different *Cannabis* smoke exposure timepoints. Ultimately, differences in the specific gestational exposure protocols (frequency and duration of dam exposure) and age of offspring during behavioral testing may account for these observed differences in behavioral outcomes.

### Amygdala gene expression following gestational *Cannabis* exposure

To the best of our knowledge, this is the first time RNAseq has been used to assess gene expression in the amygdala of rodents exposed to *Cannabis* during gestation. Analysis of these data demonstrated that *Cannabis* smoke exposure during gestation altered amygdala gene expression in a subset of male offspring during adulthood. As detailed in the results, more than half of the top 40 affected genes have known functions related to the nervous system. IPA revealed several pathways of interest, including the opioid and glutamate receptor signaling pathways. Changes in these pathways are consistent with previous reports showing altered responses to opioids and glutamate receptors in the offspring of THC-exposed rats and mice (Spano et al 2007; Frau et al 2019). Interestingly, increased expression of the preproenkephalin gene (*Penk*) was noted in the central/medial amygdala of adult mice exposed to THC while in utero (Spano et al 2007). While *Penk* levels were not significantly altered in our sample, IPA revealed that genes related to opioid signaling were increased in the amygdala samples from male offspring we collected (Fig. 8C; Supplementary Table 2B).

Some caveats related to interpretation of these data must be considered. First, tissue punches that included the bulk of the amygdala were collected, which preclude analyses of gene expression in discrete amygdala nuclei. Second, this analysis was performed on a subset of male samples as the female samples were not found to cluster when the initial PCA was performed on expression. While sex-related phenotypes have been noted following gestational *Cannabis* exposure, analysis of the present behavioral data did not reveal any sex by *Cannabis* exposure interactions in the tasks used. In addition, attempts to relate the gene expression patterns with individual offspring anxiety-like behavior in the OFT failed to account for the lack of consistently clustering in the female samples (data not shown). Given the role of the amygdala in behavioral flexibility, and reversal learning in particular, the gene networks altered in the offspring may have also contributed to the facilitated performance we observed on this task. However, as we failed to observe a sex difference here too, firm conclusions cannot be drawn. Third, while it is tempting to attribute the changes in gene expression to an enduring, direct effect of *Cannabis* exposure while the offspring were in utero, they may instead reflect compensatory changes in the amygdala from other unknown effects of *Cannabis* exposure on offspring development and behavior.

Previous research has assessed patterns of gene expression in other brain regions following gestational *Cannabis* exposure (Bara et al 2021, Scheyer et al 2019). In particular, Campolongo and colleagues (2007) found 141 genes were differentially expressed in the prefrontal cortex of male offspring of dams administered oral THC from GD15 to PND9. Networks of genes related to synaptic transmission, development and neurogenesis and myelination were altered, including several genes (*Grik3, Neurod2, Rtn4, Mobp*) that were also altered in the amygdala samples assayed in the present study. Reduced *Drd2* (DiNieri et al 2011) and increased *Penk* mRNA levels (Spano et al 2007) have been observed in the nucleus accumbens of rats exposed to THC during gestation. Others have not found differences in vesicular transporters for glutamate, GABA, and acetylcholine or SNARE proteins in cortex of the adult offspring of mice exposed to THC during gestation (Tortoriello et al 2014).

## Conclusion

Taken together, our results suggest that the repeated exposure of pregnant rat dams to high-THC *Cannabis* smoke alters some behaviors related to anxiety and behavioral flexibility in adulthood. These behavioral alterations may relate to disturbed patterns of amygdala gene expression that are also expressed in adulthood. Future research to assess the effects of other *Cannabis* products, such as high-CBD products, changes in transcript abundance in other brain regions, and other behavioral outcomes of maternal *Cannabis* exposure are now underway.

## Acknowledgements and funding sources

Funding for these experiments was obtained from the University of Saskatchewan College of Medicine and the Brain Canada Foundation to RBL and JGH. TMS was supported by a fellowship from the Saskatchewan Health Research Foundation; AJR, MAH, DLM, SNO were supported by the University of Saskatchewan College of Medicine; and TJO and SNO were supported by the Natural Sciences and Engineering Research Council of Canada.

