## Supplementary Table 1 for "Repeated exposure to high-THC *Cannabis* smoke during gestation alters sex ratio, behavior, and amygdala gene expression of Sprague Dawley rat offspring"

**Supplementary Table 1.** Statistics by figure in the main manuscript.

| **Figure/Table** | **Statistics** |
| --- | --- |
| 2B | Treatment: F(1, 140) = 1.77, p = 0.19  Time: F(6, 140) = 5.35, p < 0.0001  Interaction: F(6, 140) = 0.77, p = 0.62 |
| 2C | Treatment: F(1, 140) = 0.17, p = 0.68  Time: F(6, 140) = 122.00, p < 0.0001  Interaction: F(6, 140) = 0.67, p = 0.67 |
| 3A | t(20) = 0.344, p = 0.74 |
| 3B | t(20) = 0.12, p = 0.90  ANOVA for PND1, 7, and 14:  Treatment: F(1, 58) = 0.06, p = 0.81  Time: F(2, 58) = 1032.0, p < 0.0001  Interaction: F(2, 58) = 0.05, p = 0.95 |
| 4C | Treatment: F(1, 36) = 15.02, p = 0.0004  Sex: F(1, 36) = 1.21, p= 0.28  Interaction: F(1, 36) = 0.03, p = 0.86 |
| 4D | Treatment: F(1, 36) = 17.80, p = 0.0002  Sex: F(1, 36) = 5.74, p= 0.02  Interaction: F(1, 36) = 0.81, p = 0.37 |
| 5B | Treatment: F(1, 36) = 1.55, p = 0.22  Sex: F(1, 36) = 0.99, p= 0.33  Interaction: F(1, 36) = 0.15, p = 0.70 |
| 5C | Treatment: F(1, 36) = 0.002, p = 0.97  Sex: F(1, 36) = 1.97, p= 0.17  Interaction: F(1, 36) = 0.64, p = 0.43 |
| Table 1 – Correct Latency | Treatment: F(1, 35) = 2.23, p = 0.14  Phase: F(1, 35) = 0.33, p= 0.57  Interaction: F(1, 35) = 0.0006, p = 0.98 |
| Table 1 – Incorrect Latency | Treatment: F(1, 35) = 2.17, p = 0.15  Phase: F(1, 35) = 2.20, p= 0.15  Interaction: F(1, 35) = 0.85, p = 0.36 |
| Table 1 – Reward Latency | Treatment: F(1, 35) = 2.29, p = 0.14  Phase: F(1, 35) = 2.27, p= 0.14  Interaction: F(1, 35) = 2.01, p = 0.17 |
